# Improved *in silico* and *in vitro* methods for *E. coli* LPS outer core typing

**DOI:** 10.1101/2025.04.13.648558

**Authors:** Ellina Trofimova, Ruby P. Westerman, Paul R. Jaschke

## Abstract

The Gram-negative bacterial envelope comprises the outer membrane, periplasmic space with the peptidoglycan layer, and inner (cytoplasmic) membrane. A lipopolysaccharide (LPS) layer linked to the outer membrane is essential for survival in most species of bacteria, primarily by providing structural stability and regulating selective chemical permeability. These functions make the LPS layer a key pathogenicity determinant, protecting bacteria from host defences. At the same time, it serves as a common receptor for multiple bacteriophage orders, making it a crucial point of bacterial vulnerability. LPS outer core typing is traditionally performed using immunoblotting and PCR. With the increasing availability of sequenced genomes, PCR has emerged as the primary method for typing. This study presents a set of nine oligonucleotides designed for typing the five LPS outer core structures of *Escherichia coli*: R1, R2, R3, R4, and K-12, both *in vitro* and *in silico*. Our method was able to successfully type 99% of strains within a comprehensive dataset of 4549 complete genomes. Using these new methods, we identify previously unreported hybrid LPS structures involving R2 and K-12 outer core types and establish associations between LPS outer core types and *E. coli* phylogenetic groups, pathogenicity, O-antigen polysaccharides, and capsule types. We also introduce LPSTyper, a Python-based command-line tool that enables rapid and precise LPS outer core typing in *E. coli* genome sequences. Together, the expanded dataset of LPS outer core types and enhanced laboratory and computational tools for their detection provide new insights into LPS structural evolution and its role in *E. coli* ecology and pathogenicity. We anticipate these tools will improve the efficiency of future research and diagnostics of this important bacterial envelope structure.

**Impact Statement:** Bacterial lipopolysaccharide (LPS) layer types are important for determining both pathogenicity potential and phage susceptibility. This study presents a robust and scalable framework for precise typing of *Escherichia coli* lipopolysaccharide (LPS) outer core types, enabled by the development of nine novel oligos targeting the *waa* locus. Using the LPSTyper tool for *in silico* analysis and a simple PCR-based method for *in vitro* validation, this approach allows rapid and accurate typing across thousands of genomes, supporting large-scale monitoring of LPS diversity in both commensal and pathogenic *E. coli* populations. Our findings reveal strong associations between LPS outer core types, O-antigens, capsules, phylogenetic groups, and pathotypes, highlighting the critical role of LPS structure in virulence, immune evasion, and environmental adaptation. The discovery of hybrid and novel *waa* locus arrangements further underscores the evolutionary plasticity of LPS biosynthesis. Overall, this work provides essential tools and insights for genomic surveillance and deepens our understanding of *E. coli* surface structure variability central to host-pathogen interactions and antimicrobial resistance.

**Data Summary:** The LPSTyper source code is freely available on GitHub under the Apache 2.0 licence: https://github.com/ellinium/LPSTyper. Sequencing data for all genomes used in this study are accessible via the NCBI database under BioProject accession number PRJNA1243169. Individual genome accession numbers are listed in Table 2. All supporting data are provided in Supplementary Data Files 1 and 2.

## Introduction

The Gram-negative bacterial envelope comprises the outer membrane, periplasmic space with a peptidoglycan layer, and inner membrane. It plays a crucial role in maintaining structural integrity, acting as a selective barrier, and contributing to antibiotic resistance and host interactions. Lipopolysaccharides (LPS) are key structural components of the outer membrane in Gram-negative bacteria. They play critical roles in structural integrity and protection from external stresses, making them vital for bacterial survival. LPS contributes to antibiotic and detergent resistance by forming a permeability barrier that reduces the effectiveness of these compounds [1]. LPS is also an important target for immune recognition and response mediated by toll-like receptor 4 (TLR4), myeloid differentiation factor (MD-2), CD14, LPS-binding proteins, antimicrobial peptides, and host defence lectins[2]. In contrast to the protective effects of LPS on physical, chemical, and immune system threats, components of the LPS layer also serve as receptors for the receptor-binding proteins of diverse bacteriophage orders, representing a critical point of vulnerability [3, 4].

LPS molecules are comprised of three main parts: the lipid A region, a core polysaccharide (subdivided into inner and outer core), and the O-antigen [3]. Lipid A is a glycolipid consisting of acylated and phosphorylated β-1′-6-linked glucosamine disaccharides that anchor LPS into the membrane and are recognised by the human immune system as a potent endotoxin [5]. Glucosamines of lipid A are linked to the LPS inner core that is conserved across many bacterial species and consists of a non-repeating oligosaccharide that includes 3-deoxy-D-manno-oct-2-ulosonic acid (Kdo) and L-glycero-D-manno-heptose (Hep) residues [6]. The highly variable O-antigen, composed of one or more repeating polysaccharide units, contributes to antigenic diversity among bacterial strains, enabling *E. coli* to evade the host immune system [7]. In smooth strains, the O-antigen is attached to the outer core, creating an extended glycan barrier around the bacterium. In rough strains that lack the O-antigen, the LPS outer core represents the terminal structure exposed to the external environment.

The outer core LPS is highly variable across different Gram-negative bacterial strains [8]. This structure is primarily composed of variable hexoses and N-acetylhexosamine. Variations in the structure of the outer core can shield or expose the associated O-antigen or lipid A molecules, influencing immune evasion or activation [7]. Additionally, LPS outer core structural variability can influence bacterial susceptibility to bacteriophages. For example, certain *Siphoviridae*, *Myoviridae*, *Podoviridae* and *Microviridae* phages use components of the outer core as a receptor for attachment and infection [4]. As these structures are essential for phage infection, the LPS outer core faces conflicting selective pressures, balancing host integrity with avoiding bacteriophage recognition [4, 9].

*In E. coli*, five distinct types of outer core LPS are recognised: R1, R2, R3, R4, and K-12 [10]. The genes encoding these types are located in the *waaA* genetic locus and are organised in three operons: 1) the *waaD* operon, which is essential for the biosynthesis and transfer of L, D-heptose (synonyms: *rfaD*, *gmhD*); 2) The central *waaQ* operon containing genes necessary for the biosynthesis and modifications of the outer core; and 3) the *waaA* operon encoding Kdo transferase and additional non-LPS genes [11].

The landscape of *E. coli* outer core type distribution was previously identified in several hundred *E. coli* strains using LPS extraction followed by SDS-PAGE and immunoblotting with mouse monoclonal antibodies [12, 13] or PCR [14]. However, no *in silico* tools are currently available for *E. coli* LPS outer core typing. Accurate identification of *E. coli* LPS outer core types can provide valuable insights into immune interactions and bacterial pathogenicity, facilitate the development of targeted vaccines [15], and inform strategies to combat phage-and antibiotic-resistance.

This work describes new *in silico* and PCR-based *in vitro* methods for LPS outer core typing of any known *E. coli* strain. Using the *in silico* method, we determined the distribution of LPS outer core types across 4549 commensal and pathogenic *E. coli* strains. We validated the PCR-based method on a subset of analysed strains. Along with the identification of R1, R2, R3, R4, and K-12 types, we also identified previously uncharacterised hybrid R2 and K-12 LPS outer core gene sets. With this new dataset, we explored the associations between LPS outer core types and *E. coli* phylogenetic groups, serotypes, and capsule types, uncovering distinct correlations with specific pathogenic groups, ecological niches, and capsular and O-antigen polysaccharides. Collectively, this study significantly enhances our understanding of the association of *E. coli* LPS outer core types, virulence, and adaptability, while also providing user-friendly tools for further investigation of these associations.

## Methods

### Bacterial strains

*E. coli* reference collection (ECOR) from the Yehl Lab (Miami University) was verified with multiple-locus variable-number tandem-repeat analysis using multiplex PCR[16]. *E. coli* ROAR340, DSM13127, ROAR375, C4741, M1402 were kindly provided by Olivier Tenaillon (Institut Cochin, Université Paris Cité). *E. coli* NCTC122 was from the National Collection of Type Cultures-UK Health Security Agency.

### Bacterial genomic data

A total of 4549 complete *E. coli* genomes were retrieved from the NCBI Reference Sequence database in March 2024. The dataset included commensal and pathogenic strains. The complete list of genomes is provided in Supplementary Table S1. Pathogenic strains were subtyped based on the keywords related to pathogenicity in the metadata provided with the genomes from the columns ‘Assembly BioSample Description Title’, ‘Assembly Accession’, ‘Assembly BioSample Description Comment’, ‘Assembly BioProject Lineage Title’, ‘Assembly BioSample Attribute Name’, ‘Assembly BioSample Attribute Value’. All genomic data were downloaded in FASTA nucleotide format and re-annotated with Prokka v1.13 [17] using the following parameter values: ‘genus’ - Escherichia, ‘species’ - coli, ‘kingdom’ - Bacteria, ‘gcode’ - 11, ‘usegenus’, and ‘proteins’ referring to a MG1655 FASTA amino acid file (GenBank accession ID U00096.3[18]) used for the EcoCyc database [19].

### LPS outer core identification

Genomic data was parsed using custom Python scripts to obtain *waa* operon gene sequences from *waaD* (*rfaD)* to *coaD*, including nucleotide and amino acid sequences. The sequences and their order were compared to previously typed strains and LPS operon gene order described in the literature [11, 14, 20] and aligned using the online Clustal Omega service ver. 2.1 [21]. Primers for the LPS outer core were identified semi-manually using Geneious Prime® 2024.0.7. For genes in question, annotations were compared with the original GenBank data, searched using NCBI Nucleotide and Protein Blast online services, Uniprot Blast [22], and aligned with the known genes.

### In vitro LPS outer core typing

PCR was performed on BIO-RAD T100™ Thermal Cycler using OneTaq® Hot Start DNA Polymerase (NEB) according to the manufacturer’s protocol. The program included an initial denaturation at 94°C for 30 seconds, followed by 30 cycles of denaturation at 94°C for 30 seconds, annealing at 49°C for 30 seconds, extension at 68°C for 90 seconds, and a final extension at 68°C for 5 minutes. The PCR products were separated on the 1% agarose gel containing thiazole orange (0.1 μl/ml) for 40 min.

The same PCR program was applied for PCR with all LPS primers combined in the same mixture (multiplex PCR), with individual primers used at a final concentration of 0.2 μM (Table 1). The template DNA was prepared as described previously, with one μl used per a 10 μl reaction [16].

**Table 1.**
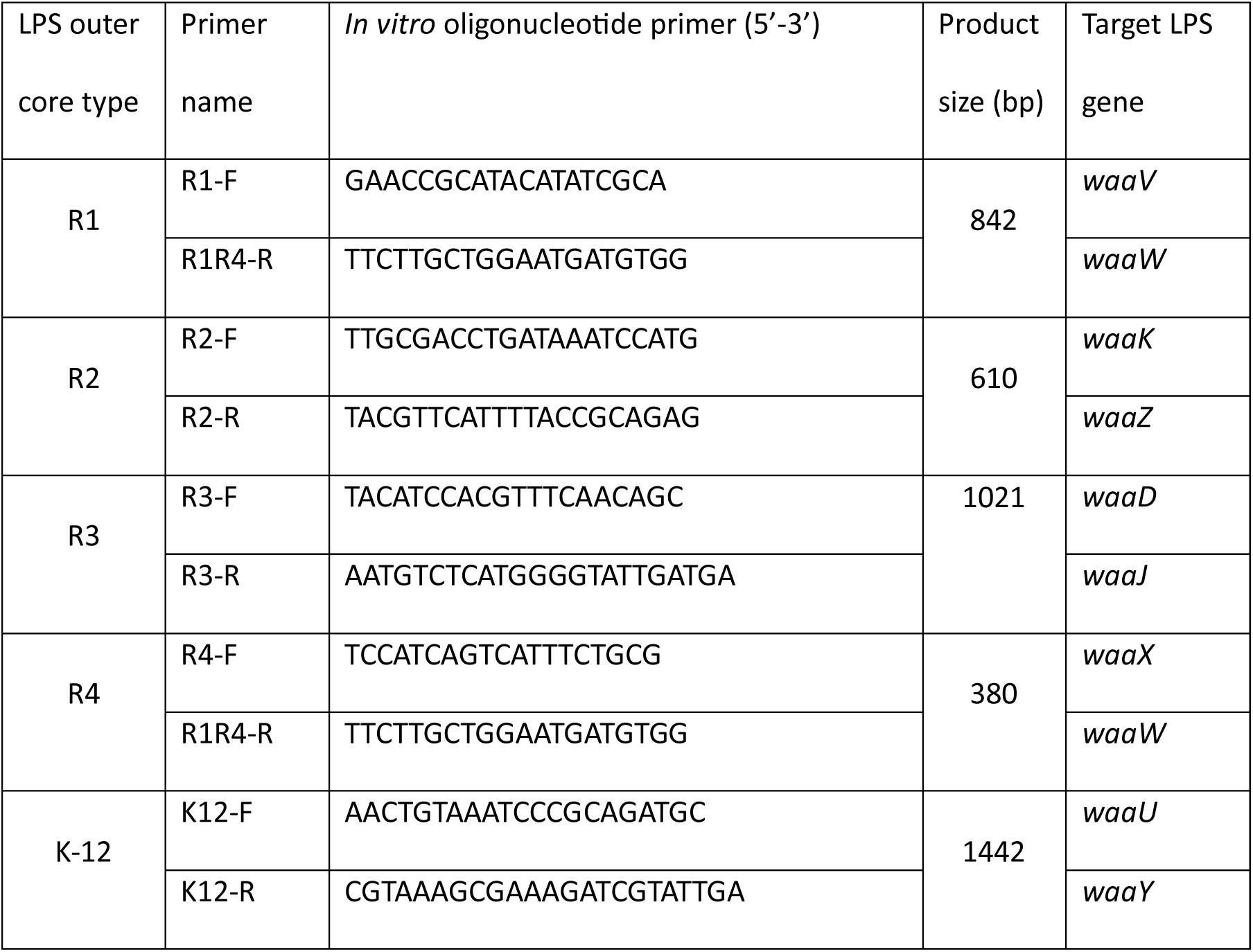
Oligonucleotide primers for LPS outer core typing.

### Determination of E. coli phylogenetic groups, serotypes and capsules

*E. coli* phylogenetic groups were determined using a standalone version of the Clermont phylotyping tool ClermonTyping v23.06 [23]. Serotypes were identified with a command-line version of SerotypeFinder 2.0 [24], and capsules were typed with VirulenceFinder 2.0 [25]. The capsule was identified if both *kpsE* and *kpsM* genes were present.

### Sequencing

Genomic DNA from *E. coli* ROAR340, DSM13127, ROAR375, C4741, M1402 were extracted using Qiagen DNeasy Blood & Tissue Kit (Gram-negative bacterial protocol), sequenced at ithree institute (University of Technology, Sydney) on Illumina MiSeq in 2×150bp mode and further assembled and annotated as described previously [26]. PCR products for R1, R2, R3, R4 and K-12 LPS core types were purified using Monarch® DNA clean-up kit (NEB) and sequenced using Plasmidsaurus service ‘Standard Purified Linear PCR’. The bacterial genomes were deposited in NCBI BioProject PRJNA1243169.

## Results

Surveying current methods for identifying LPS outer core types in *E. coli*, we found that existing approaches relied solely on wet lab methods, with a noticeable gap in computational tools. We then attempted to adapt previously published PCR oligonucleotides [14] for *in silico* identification of the LPS outer core genes. Using an *E. coli* dataset comprising 4549 genomes, including commensal and pathogenic strains, we found that only 46.81% of the genomes could be LPS outer core typed using these oligonucleotides for *in silico* detection. Although PCR can tolerate single-point mismatches in primer binding sites *in vitro*, *in silico* typing requires identifying all mutations in the primer-binding region beforehand. To address this issue, we aimed to create primers for LPS outer core typing that could be used both *in silico* and *in vitro*.

Using aligned nucleotide sequences of the genes within the *waa* locus of *E. coli* strains with known LPS outer core types [20] we designed nine PCR oligonucleotides (Table 1) to uniquely identify genes diagnostic for R1, R2, R3, R4 and K-12 LPS outer core types. We iteratively refined the oligos *in silico* on the genomes of 4549 *E. coli* strains until 99% of the LPS outer core types were identified.

The final oligo sequences (Table 1) were bound to a 842-nt region in R1 types between UDP-galactose-(Galactosyl) LPS α-1,2-galactosyltransferase (*waaW*) and β-1,3-glucosyltransferase (*waaV*), a 610 bp region between the gene involved in KdoIII attachment during LPS core biosynthesis (*waaZ*) and the lipopolysaccharide 1,2-N-acetylglucosamine transferase (*waaK*) for R2, and a 1021 bp segment between the UDP-glucose: (galactosyl) LPS α-1,2-glucosyltransferase (*waaJ*) and ADP-L-glycero-D-mannoheptose 6-epimerase (*waaD*) genes for R3 (Figure 1). For R4 outer core typing, the same oligo used for identifying the *waaW* gene in R1 types was paired with a unique sequence in β-1,4-galactosyltransferase (*waaX*), yielding a 380 bp product. For the K-12 type, a 1442-nt region was identified between lipopolysaccharide core heptose (II) kinase *waaY* and the ADP-L-glycero-β-D-manno-heptose:LPS heptosyltransferase IV (*waaU*).

**Figure 1:**
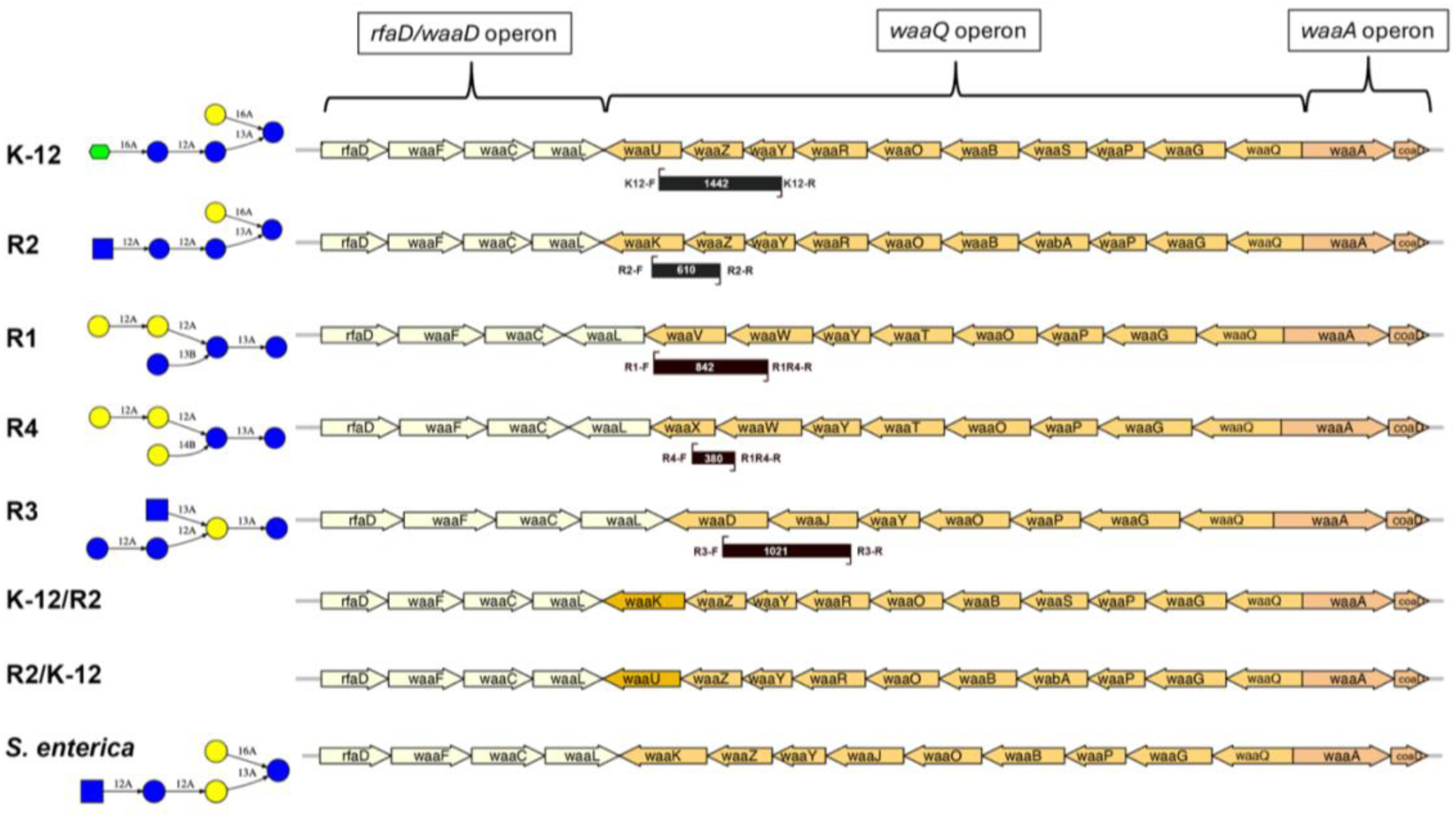
Genomic organisation of *E. coli* LPS operons and corresponding glycan structures across eight outer core types. The outer core types R1-R4 and K-12 represent distinct *E. coli* variants. K-12/R2 and R2/K-12 illustrate hybrids of *E. coli* K-12 and R2 types. Genes are aligned to facilitate the comparison of outer core architectures. Gene *waaG* overlaps with *waaQ* and *waaP* in all *E. coli* core types [27]. *Salmonella enterica* Typhimurium LPS genes are included for comparison with R2 and K-12 genes. A complete gene annotation is provided in Table S9. Monosaccharides are depicted using the Symbol Nomenclature for Glycans (SNFG) [28]: yellow circles represent galactose (Gal), blue circles represent glucose (Glc), blue squares represent N-acetylglucosamine (GlcNAc), green flat hexagon represents L-Glycero-D-Manno-Heptose (LDmanHep). Glycosidic linkages are indicated by black lines, with numbers specifying the bond positions and anomeric configurations - α1→6 (16A), α1→2 (12A), α1→3 (13A), and β1→4 (14B).

### In vitro determination of E. coli LPS outer core types using PCR

We evaluated the primers *in vitro* using *E. coli* strains NCTC122 (R1 type), ECOR11 (R2), ECOR09 (R3), ECOR04 (R4), and MG1655 (K-12) (Table 2). The PCR products were of the expected sizes (Figure 2), and sequencing confirmed that the amplified products matched the correct target sequences. We repeated the test with a multiplex primer set combining all nine primers and had the same results. We next performed PCR on additional strains, including those previously typed in Michel *et al*. [20] and two ECOR strains. In all cases, the results were consistent with those reported in the original publications (Table 2), showcasing the ease and utility of our new PCR primers for LPS outer core typing.

**Table 2.**
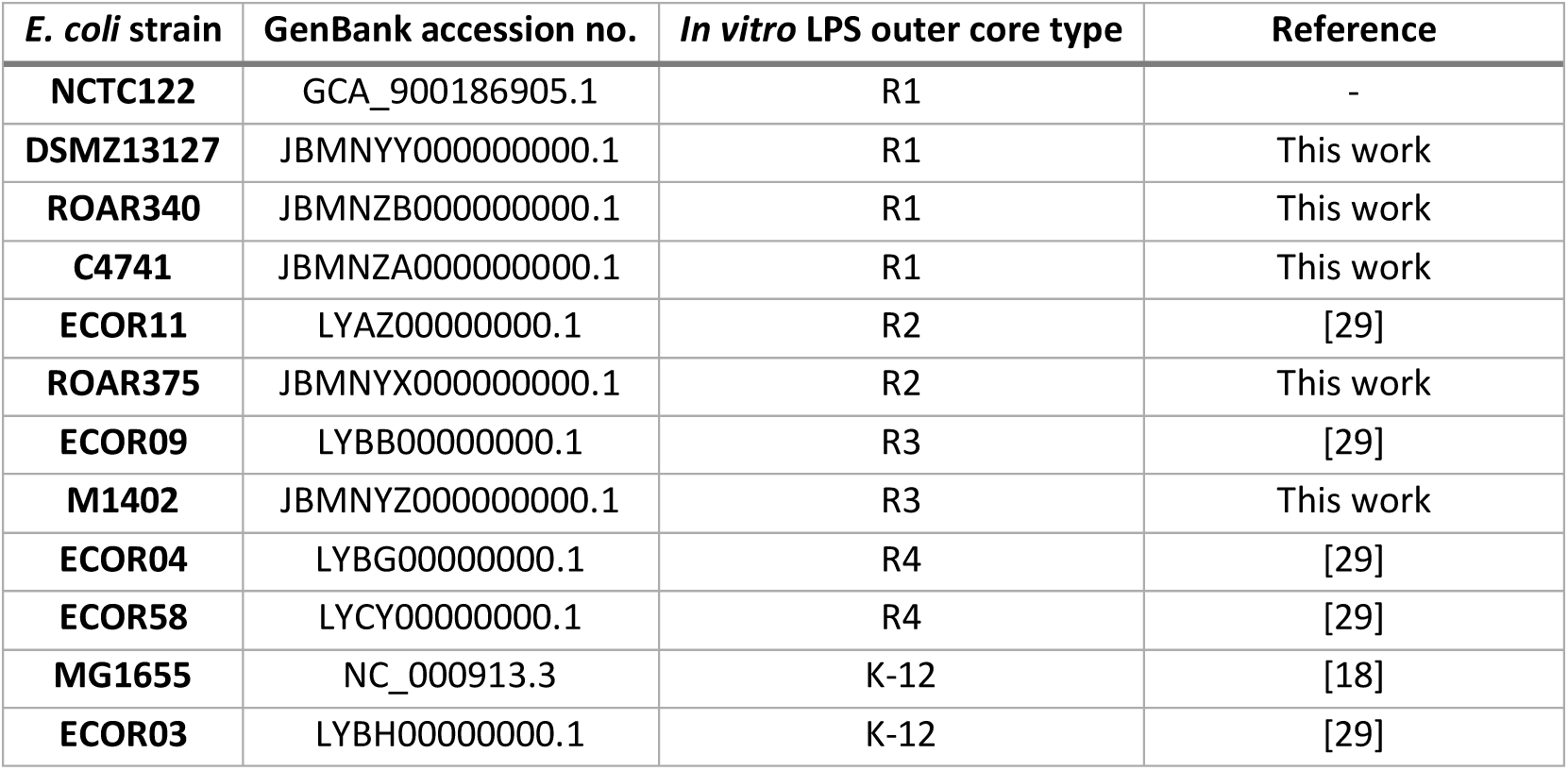
Reference *E. coli* strains used in *in vitro* PCR.

**Figure 2.**
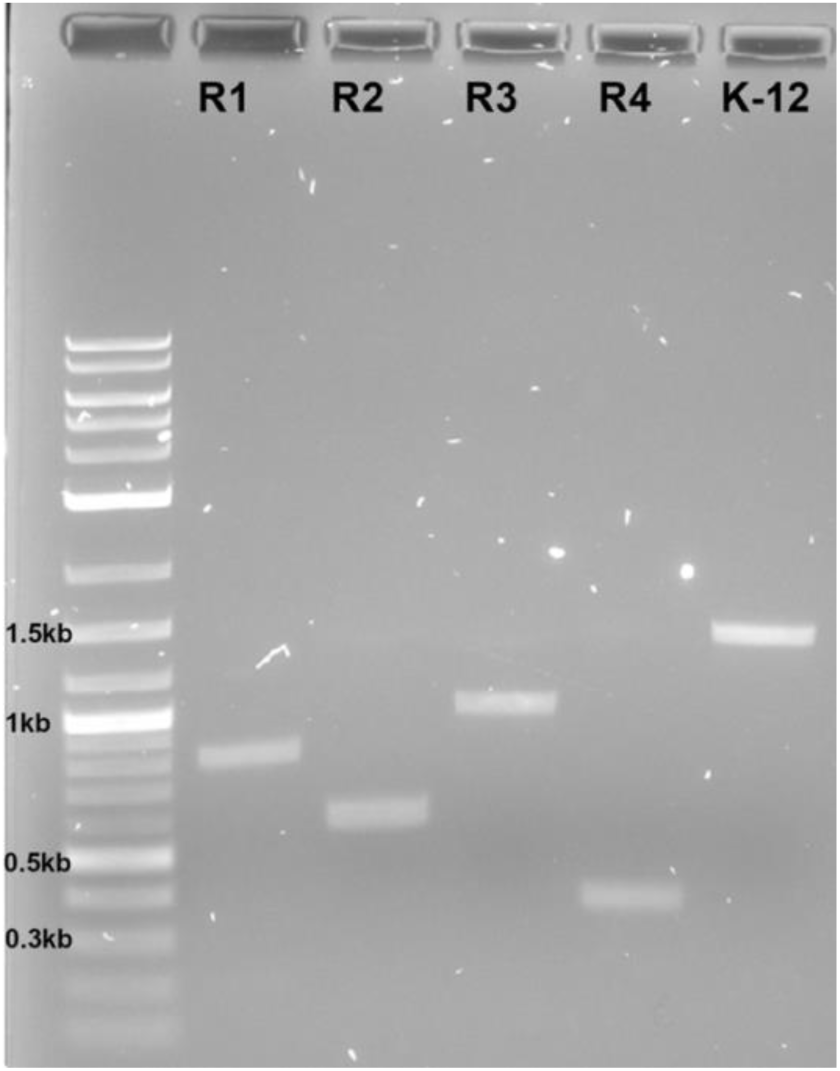
PCR products amplified from five different *E. coli* LPS outer core types separated on a 1% agarose gel. The following strains were used for the typing: *E. coli* NCTC122 (R1), ECOR11 (R2), ECOR09 (R3), ECOR04 (R4), and MG1655 (K-12).

### In silico determination of E. coli LPS outer core types

We next developed LPSTyper, a command-line Python tool designed to automatically identify LPS outer core types from nucleotide sequences (https://github.com/ellinium/LPSTyper). The tool accepts a single input parameter: the full path to a directory containing one or more files in FASTA nucleotide sequence format (e.g., FASTA, FNA). It scans these files using our *in silico* LPS outer core oligos to identify sequences matching with forward and reverse primers and then uses presence/absence patterns to determine the corresponding LPS core type—R1, R2, R3, R4, or K-12. LPSTyper supports recursive processing of subdirectories within the input directory. For files containing contigs, the LPS *waa* operon genes must either reside on the same contig, or the contig order must follow the sequential order of the genes in the *waa* operon in either direction. The tool does not work with FNA or FASTA files containing coding sequences and does not correct genome assembly errors.

### ECOR collection verification

We used LPSTyper to identify the LPS outer core types in the *E. coli* reference collection (ECOR), which returned results for all strains except ECOR46 (Table 3). We could not initially identify the ECOR46 LPS outer core type from its nucleotide sequence using *in silico* primers due to the genome sequence not being closed as a single contig, and the LPS genes being on different contigs and out of order. Additionally, we discovered two copies of O-antigen ligase *waaL*: one matching R1 *waaL* and another matching R4 *waaL*. Moreover, both R1 *waaV* and R4 *waaX* genes were upstream of corresponding *waaL* genes. We also found a transposase gene *insO* upstream of *waaY*, suggesting that LPS gene order was changed and both R1 and R4 genes appeared after a transposition event. This unusual genome configuration could be a reason for the R1 typing results in Amor *et al*. [14] and R4 in Gaborieau *et al*. [30] We PCR-typed ECOR46 *in vitro* using the primers developed in this work (Table 1) and obtained an amplification product matching only the R4 core type, not R1.

**Table 3.** ECOR collection LPS outer core types.

| ECOR strain | GenBank accession no. | *In silico* LPS outer core type prediction<sup>1</sup> |
| --- | --- | --- |
| 1 | LYBJ000000000.1 | R2 |
| 2 | LYBI000000000.1 | K-12 |
| 3 | LYBH000000000.1 | K-12 |
| 4 | LYBG000000000.1 | R4 |
| 5 | LYBF000000000.1 | R2 |
| 6 | LYBE000000000.1 | R3 |
| 7 | LYBD000000000.1 | K-12 |
| 8 | LYBC000000000.1 | R2 |
| 9 | LYBB000000000.1 | R3 |
| 10 | LYBA000000000.1 | R2 |
| 11 | LYAZ000000000.1 | R2 |
| 12 | LYAY000000000.1 | R2 |
| 13 | LYAX000000000.1 | K-12 |
| 14 | LYAW000000000.1 | K-12 |
| 15 | LYAV000000000.1 | R1 |
| 16 | LYAU000000000.1 | R1 |
| 17 | LYAT000000000.1 | R1 |
| 18 | LYAS000000000.1 | R1 |
| 19 | LYAR000000000.1 | R1 |
| 20 | LYAQ000000000.1 | R1 |
| 21 | LYAP000000000.1 | R1 |
| 22 | LYAO000000000.1 | R1 |
| 23 | LYAN000000000.1 | R1 |
| 24 | LYAM000000000.1 | R1 |
| 25 | LYAL000000000.1 | R2 |
| 26 | LYAK000000000.1 | R3 |
| 27 | LYAJ000000000.1 | R3 |
| 28 | LYAI000000000.1 | R3 |
| 29 | LYAH000000000.1 | R1 |
| 30 | LYAG000000000.1 | R1 |
| 31 | LYAF000000000.1 | R2 |
| 32 | LYAE000000000.1 | R1 |
| 33 | LYAD000000000.1 | R1 |
| 34 | LYAC00000000.1 | R1 |
| 35 | LYAB00000000.1 | R1 |
| 36 | LYBO00000000.1 | R1 |
| 37 | LYAA00000000.1 | R3 |
| 38 | LXZZ00000000.1 | R1 |
| 39 | LYCJ00000000.1 | R1 |
| 40 | LYCI00000000.1 | R1 |
| 41 | LYCH00000000.1 | R1 |
| 42 | LYCG00000000.1 | R1 |
| 43 | LYCF00000000.1 | R2 |
| 44 | LYCE00000000.1 | R3 |
| 45 | LYCD00000000.1 | R3 |
| 46 | LYCC00000000.1 | R1/R4 hybrid |
| 47 | LYCB00000000.1 | R3 |
| 48 | LYCA00000000.1 | R4 |
| 49 | LYBZ00000000.1 | R4 |
| 50 | LYBY00000000.1 | R4 |
| 51 | LYYB00000000.1 | R1 |
| 52 | LYCT00000000.1 | R1 |
| 53 | LYCU00000000.1 | R1 |
| 54 | LYCV00000000.1 | R1 |
| 55 | LYCW00000000.1 | R1 |
| 56 | LYDK00000000.1 | R1 |
| 57 | LYCX00000000.1 | R1 |
| 58 | LYCY00000000.1 | R4 |
| 59 | LYCZ00000000.1 | R1 |
| 60 | LYDA00000000.1 | R1 |
| 61 | LYDB00000000.1 | R1 |
| 62 | LYDL00000000.1 | R1 |
| 63 | LYDM00000000.1 | R1 |
| 64 | LYDC00000000.1 | R1 |
| 65 | LYDD00000000.1 | R1 |
| 66 | LYDE00000000.1 | R1 |
| 67 | LYDN00000000.1 | R1 |
| 68 | LYDF00000000.1 | R1 |
| 69 | LYDG00000000.1 | R1 |
| 70 | LYDH00000000.1 | R1 |
| 71 | LYDI00000000.1 | R1 |
| 72 | LYDJ00000000.1 | R3 |

<sup>1</sup>Results from *in silico* experiments performed in this work – see Materials and Methods. The comparison with Amor *et al.* [14] and Gaborieau *et al.* [30] is provided in Supplementary Table S2.

Our *in silico* results for the remaining ECOR strains revealed discrepancies with Amor *et al*. [14] for strains ECOR07, ECOR43, ECOR44, ECOR46, ECOR47-ECOR50, ECOR58 and ECOR72, and Gaborieau *et al*. [30] for strains ECOR43 and ECOR46 (Supplementary Table S2). For ECOR07, both our tool and Gaborieau *et al*. results showed K-12, while Amor *et al*. typed this strain as R1. ECOR43 had R4 type in Amor *et al*. and was non-typed in Gaborieau *et al*. With LPSTyper and additional manual verification of the LPS gene sequences, we identified it as R2. ECOR44, ECOR47, and ECOR72 all had R3 type, while ECOR48, ECOR49, ECOR50, and ECOR58 had R4 type in our data and Gaborieau *et al.* By contrast, Amor *et al*., identified all those strains as R1 type except ECOR58, which had R3 type (Supplementary Table S2). Overall, these contrasting results may be attributed to the source of the ECOR collection and the level of cross-contamination, as previously documented [16].

### Hybrid LPS and non-typed structures

Next, we looked closer at *E. coli* strains that did not return an LPS outer core type identification from LPSTyper. Manual genome analysis revealed several genomes that should have been identified as R1 or R4 types but had single-point mutations in the gene locations where the *in silico* primers bound, thus preventing their identification. To address this, we included additional *in silico* primers (Table S10) in our command-line typing tool LPSTyper. We took an approach of including primers that identified >99% of samples we analysed while avoiding adding additional primers for every possible variation in the LPS genes targeted by PCR as they are highly conserved within each LPS outer core type. When required, new primers can be incorporated into the LPSTyper tool in the future to account for any newly discovered mutations in these target genes.

### R2/K-12 hybrids

While manually identifying the LPS outer core type of *E. coli* strains that could not be typed using LPSTyper, we discovered several strains that possessed gene sets from both R2 and K-12 types (Table S11). Eight strains had genes identical to K-12 type but with *waaU* gene (ADP-L-glycero-β-D-manno-heptose:LPS heptosyltransferase IV) replaced by *waaK* (lipopolysaccharide 1,2-N-acetylglucosamine transferase) from R2 type with 90% nucleotide identity. In these strains, the *waaL* gene was also 100% identical to *waaL* of K-12, and *waaS* was 93.25% identical to K-12 *waaS* (Figure 1).

Two additional genomes, GCA_036870985.1 and GCA_012711215.2, were also identified with gene sequence arrangements identical to the K-12 core genes but with a putative transposase, *yhhI*, inserted between *waaZ* and *waaY*. Furthermore, nucleotide identity analysis revealed that *waaL* shared 92.86% identity and *waaK* 89.94% identity with the corresponding *waaL* and *waaK* genes of R2 type, while *waaS* exhibited 93.16% identity to the *waaS* gene of K-12. Moreover, we identified one strain (GCA_013122565.1) with an R2 core gene arrangement but with *waaK* replaced by K-12 *waaU*, and *waaL* and *wabA* genes matching R2 gene sequences (Figure 1). These findings suggest transposition or horizontal gene transfer events have occurred in *E. coli* strains containing R2 and K-12 outer core structures.

We also identified two strains with duplicated LPS genes. *E. coli* MSB1_3C (GCA_904863165.1) contained a gene sequence resembling the R2 structure but included two truncated copies of each of the *waaK* and *waaQ* genes. Similarly, *E. coli* O158:H23 (GCA_018986795.1) exhibited a K-12-like structure but with two *waaS* genes and an additional short peptide sequence between *waaA* and *waaQ*. These findings may represent either genome assembly artefacts or genuine gene rearrangement events. There was also one strain (GCA_013892175.1) following K-12 structure with an additional short 96-nucleotide sequence between *waaA* and *waaQ* genes matching GCA_018986795.1 strain and encoding an uncharacterised protein (UniProt ID A0A485J7Q0).

### Escherichia marmotae

Interestingly, we found two genomes (GCA_013898955.1, GCA_013898975.1) that were initially classified as *E. coli* and possessed K-12 LPS gene sequence order. However, nucleotide alignment showed lower identity to K-12: *waaL* (48.93%), *waaY* (90.27%), *waaU* (82.21%), *waaS* (91.25%). We later found that those strains were re-classified as *Escherichia mormotae* [31], suggesting LPS biosynthetic pathways share a close genetic relationship between *E. coli* K-12 and *E. mormotae*.

### Distribution of LPS outer core types within commensal and pathogenic E. coli

To investigate the association between *E. coli* LPS outer core types and pathogenicity, we categorised the dataset into several pathogenic and non-pathogenic groups, each containing more than 20 strains. Pathogenic groups included enterohaemorrhagic *E. coli* - EHEC (n =187), enterotoxigenic *E. coli* - ETEC (n=44), avian pathogenic *E. coli* - APEC (n=23), neonatal meningitis *E. coli* - NMEC (n=27), uropathogenic *E. coli* - UPEC (n=369), and other non-typed extraintestinal pathogenic *E. coli* - ExPEC (n=21). Additionally, 1394 strains were classified as pathogenic without subtype designation. The remaining strains consisted of just under half the dataset (n=2452) and were identified as commensal strains. Strains with unidentified LPS core types (n=44) were excluded from the analysis.

Our findings indicated that the R1 LPS outer core type was the most prevalent among all *E. coli* strains analysed, accounting for 35.38% (Figure 3A). It was followed by R3 (25.79%), K-12 (18.38%), R4 (12.3%), and R2 type (8.15%) as the least common. Commensal *E. coli* strains showed a similar distribution as the whole dataset, with the R1 LPS type being most prevalent (36.9%), followed by R3 (23.4%), and R2 being the least common (8.7%) (Figure 3B).

**Figure 3.**
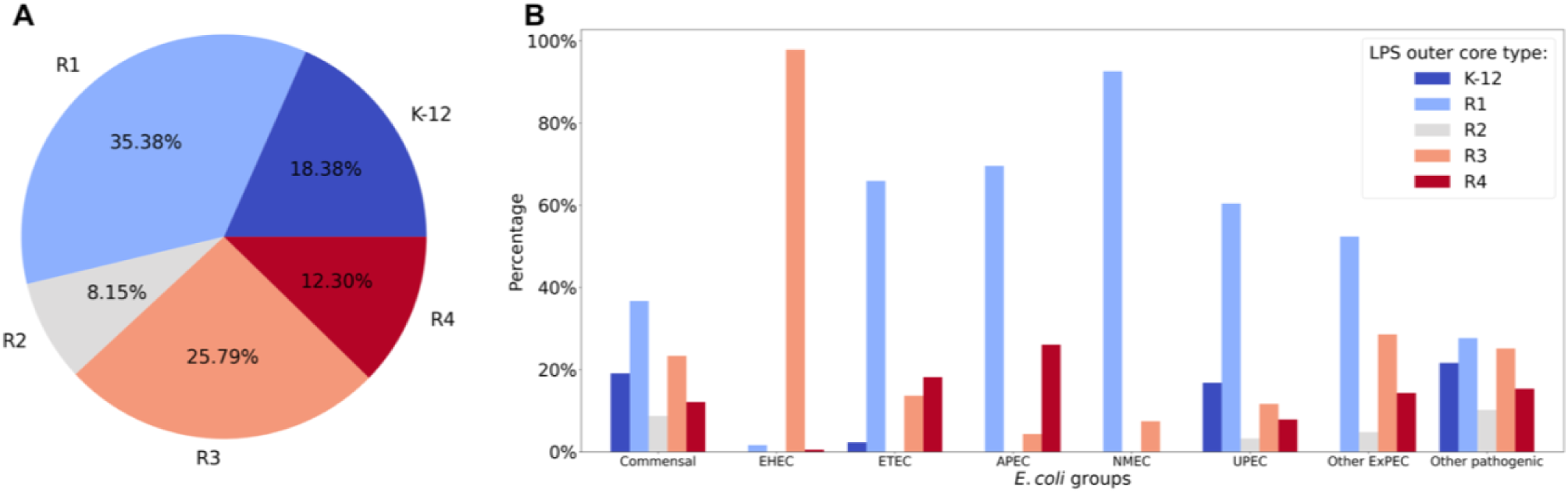
Distribution of *E. coli* LPS outer core types within pathogenic and commensal groups. (A) Distributions of LPS outer core types within 4505 commensal and pathogenic *E. coli* strains with identified LPS outer core types. (B) Distribution of LPS outer core types within different pathogenic subgroups of *E. coli* and commensal strains. The numerical data for the figure is available in Supplementary Table S3.

By contrast, the distribution of LPS outer core types across different *E. coli* pathogenic types revealed significant variations, underscoring associations between specific LPS structures and *E. coli* pathotypes (Figure 3B). EHEC strains were predominantly associated with the R3 core type (97.9%), with the remaining ∼2.1% composed of R1 and R4. This group is primarily represented by *E. coli* O157:H7 strains in databases, which may constrain the observed LPS outer core genetic variability. ETEC strains were predominantly associated with the R1 LPS outer core type (65.9%), with moderate representation of R4 (18.2%) and R3 (13.6%). No ETEC strains were associated with the R2 type.

Within the ExPEC pathotype, NMEC strains showed a near-exclusive association with the R1 LPS type (92.6%), with minor representation in R3 (7.4%) and no presence in other LPS types (Figure 3B). Likewise, APEC strains were primarily associated with the R1 core (69.57%), with a smaller proportion linked to the R4 type (26.09%). The other ExPEC group, UPEC, was also dominated by R1 LPS type (60.4%) but exhibited the remaining LPS outer core types: K-12 (16.8%), R3 (11.6%), R4 (7.9%), and R2 (3.2%). ExPEC strains that could not be classified into a subtype were dominated by R1 type (52.4 %) but appeared to match the overall LPS type pattern of UPEC strains rather than NMEC strains. However, the K-12 LPS type was conspicuously absent from this group, suggesting the presence of an additional unclassified subtype within ExPEC (Figure 3B).

Pathogenic strains that were not split into subgroups displayed a more even distribution, with moderate prevalence in R1 (27.7%), R3 (25.2%), K-12 (21.6%), and R4 (15.3%), and a lower association with R2 (10.2%), likely a result of the mixed nature of the grouping.

Across all *E. coli* populations, the R1 LPS outer core structure was the most dominant except in EHEC strains, while the R2 type was the least common and entirely absent in EHEC, NMEC, and ETEC strains. The K-12 LPS structure was also lacking in EHEC and NMEC strains. These results strongly suggest that *E. coli* pathogroups show selection for specific LPS outer core types, indicating a role for LPS structure in influencing bacterial colonisation, adaptation, or pathogenicity.

### Distribution of LPS outer core types within E. coli phylogenetic groups

The established phylogenetic groups of *E. coli* (A, B1, B2, C, D, E, F, and G) represent bacteria occupying different ecological niches with sets of similar characteristics, such as antibiotic resistance, metabolic capacity, and pathogenicity [32, 33]. Using the LPSTyper results, we compared the LPS outer core distribution across these eight major *E. coli* phylogroups.

Commensal phylogroup A (n=1352) showed the most balanced distribution among the outer core types, with R1, R2 and K-12 present in similar proportions at 28.8%, 25.5% and 28.6%, respectively (Figure 4). Phylogroup B1 (n=891) was highly dominated by R1 (42.5%) and R3 (49.7%), while B2 (n=696) was dominated by K-12 (43.6%) and R1 (53.4%). By contrast, phylogroup C (n=165) was almost exclusively associated with R1 (99.4%). Phylogroup D (n=380) exhibited the highest proportions for R4 (60.9%) and R3 (28.9%), whereas phylogroup E (n=436), which accounted for more than half of the pathogenic strains, was predominantly associated with the R3 core type (84.3%). Phylogroup F (n=128) showed a high proportion of R4 (51.4%) and R1 (45.7%), whereas phylogroup G (n=49) had the highest proportion of R4 (84.9%) among all groups. These results clearly demonstrate that each phylogroup has a distinct pattern of LPS outer core type distribution and provide the first report, to our knowledge, on LPS outer core types for phylogroup G.

**Figure 4.**
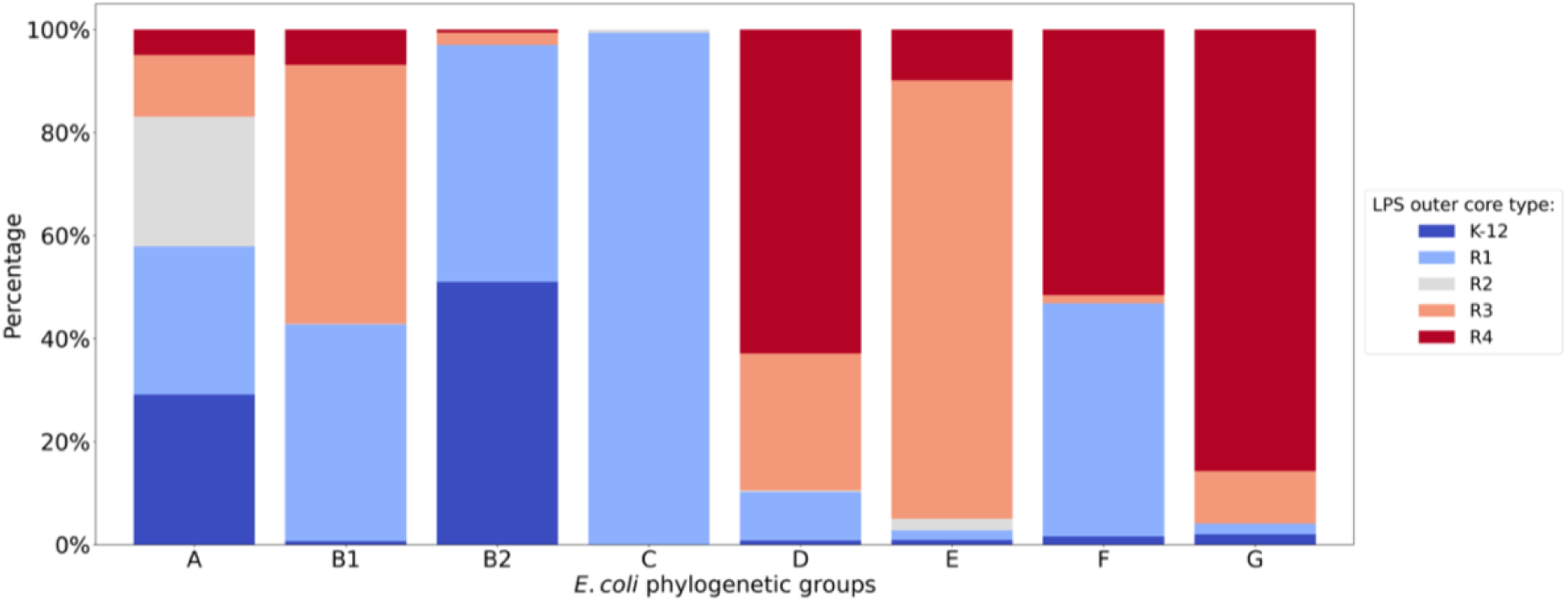
Distribution of LPS outer core types within *E. coli* phylogenetic groups. The numerical data for the figure is available in Supplementary Table S4.

### Distribution of LPS outer core types within E. coli serogroups

During LPS assembly, the O-antigen is covalently attached to the outermost hexose of the LPS outer core by the O-antigen ligase WaaL. Due to the structural and chemical requirements of this reaction, we hypothesised that there might be an association between O-antigens and LPS outer core types. To investigate this, we analysed the distribution of O-antigens among *E. coli* strains with LPS outer core types determined by LPSTyper and examined the pairwise frequency of each core type with predicted O-antigens across our *E. coli* genome dataset.

We found that all O-antigens were strongly associated with only one LPS outer core type except for O1, O5, O83, O128, and O141, which were associated with multiple core types (Table S5). The O128 serotype frequently co-occurred with R1 (35.0%) and R3 (30.0%), while O83 was evenly distributed between R1 (46.1%) and R4 (53.8%) types. Similarly, O141 showed 42.9% co-occurrence with both R2 and R3. By contrast, the O5 serotype displayed a broader distribution, associating with K-12 (34.3%), R1 (28.6%), and R3 (34.3%) (Table S5). Our analysis revealed that a total of 136 out of 197 (69.0%) predicted O-antigen types co-occurred at least once with the R1 outer core type, 64.0% with R3, 38.1% with R4, 31.5% with R2 and 17.8% with the K-12 outer core type. By contrast, K-12 and R2 outer core types were found at low frequencies or were absent for many O-antigen types.

To measure the strength of association between specific LPS outer core types and O-antigen types, we calculated Cramér’s V statistics with a bias correction to quantify the effect size [34]. Although we found 197 serotypes in our data, including mixed types, only 29 serotypes reached abundances necessary to calculate the association with LPS core types (Figure 5A, Table S6). We identified a statistically significant and very strong association between O16 O-antigen - one of the most common EHEC-associated O-types [35] - and the K-12 LPS outer core type (Cramér’s V=0.62). Similarly, O25, a serogroup frequently found in ETEC strains [35], was also associated with K-12 (0.54), while O157 showed a very strong association with R3 (0.67). Meanwhile, the R2 type was strongly associated with the O89 serogroup (0.79), which has recently been shown to be a rough strain serotype that lacks any expressed O-antigen [7]. We also observed moderate associations between O1 (0.21) - O-type primarily linked to bloodstream and urinary tract infections [36], O102 (0.36) and O86 (0.29) serotypes with R4, as well as O157 (0.26), O6 (0.26), O8 (0.27), O16 (0.25), and O25 (0.21) serotypes with R1.

**Figure 5.**
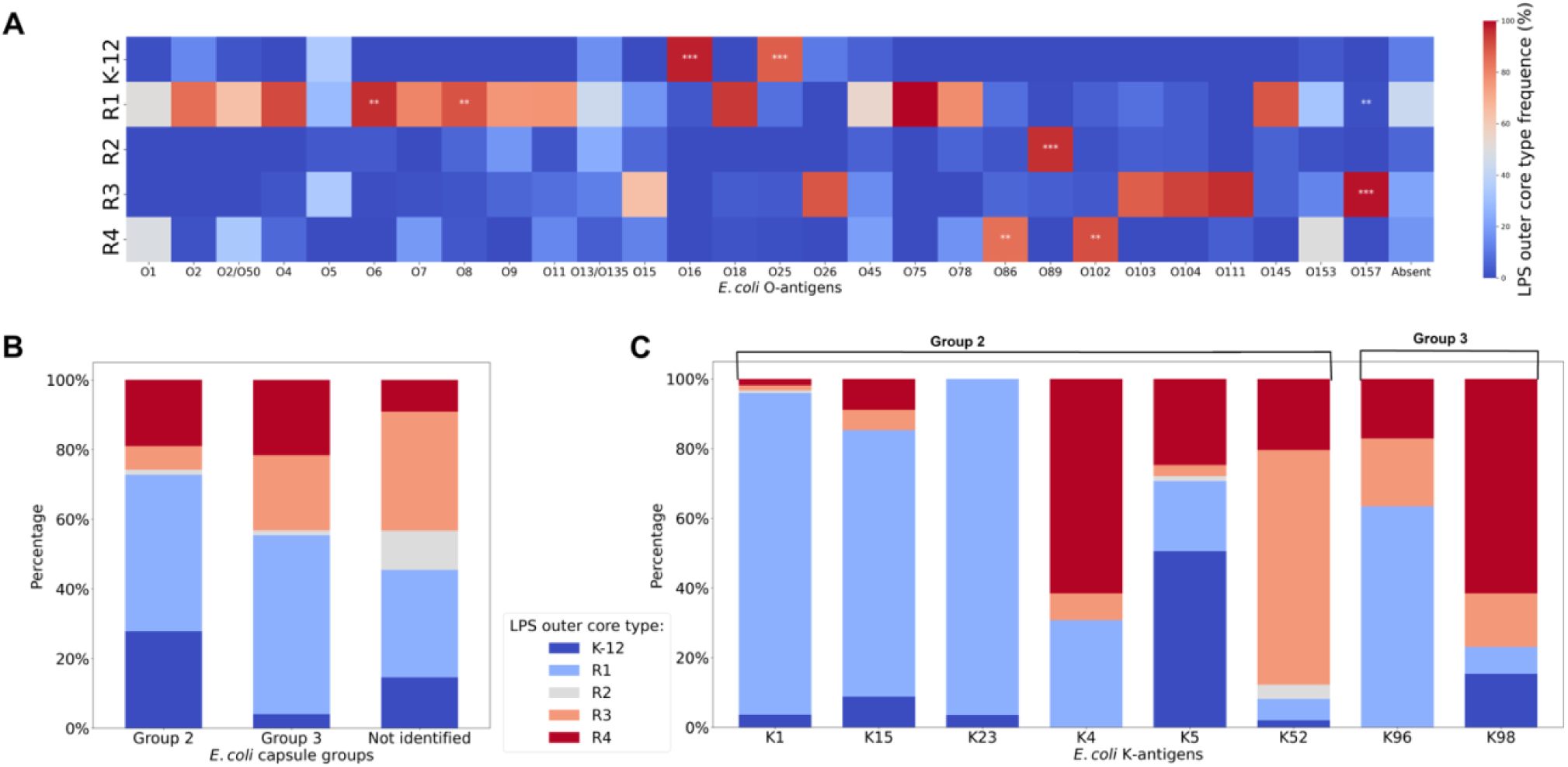
Distribution of *E. coli* LPS outer core types within select O-antigens, capsule groups and K-antigens. (A) A heatmap of associations between *E. coli* LPS outer core types and serotypes with calculated Cramér’s V. Cramér’s V association effect is denoted on the heatmaps as *** very strong association (effect size, >0.6) or ** moderate association (0.2 < effect size ≤ 0.6). Weak associations are not shown. (B) *E. coli* LPS outer core type distribution within capsule groups. (C) *E. coli* LPS outer core type distribution within specific capsule K-antigens. The numerical data for the figure is available in Supplementary Tables S6, S7, and S8.

These findings suggest a potential co-evolutionary relationship between O-antigens and LPS outer core types likely driven by selective pressures ensuring structural compatibility, stability, and functional synergy [37].

### Distribution of LPS outer core types within E. coli capsule types

The *E. coli* capsule - which is typically represented by K-antigens [38] - along with the LPS core and O-antigen polysaccharides, are structurally and functionally interconnected, with shared biosynthetic pathways and coordinated assembly ensuring outer membrane stability and protection [39]. This relationship enhances immune evasion, as the capsule masks O-antigens, while the LPS core provides structural support for capsule and O-antigen attachment and bacterial virulence [2, 38]. The presence of the capsule can also differentiate *E. coli* strains based on their susceptibility to phage predation. In some instances, the capsule can protect *E. coli* from phage infection by obstructing the LPS receptor [40]. Conversely, certain phage species can use the capsule as an initial receptor, binding and degrading a hole through it. This process exposes the LPS outer core, which is then used as the terminal receptor, allowing phage entry into the bacterial cytoplasm [40].

We analysed the distribution of LPS core types within strains with group 2 and group 3 capsules, as well as *E. coli* lacking capsules (Figure 5B). For strains with group 2 (n=1358) and 3 (n=74) capsules, the R1 outer core type was the most prevalent, accounting for 45.0% and 51.3%, respectively. The R4 core type showed similar frequency distribution in both groups (19.0% and 21.6%), whereas the K-12 core type was significantly more common in group 2 than in group 3 (27.8% vs. 4.0%). Conversely, the R3 core was more frequent in group 3 (21.6%) compared to group 2 (6.8%).

In strains lacking capsules (or with unidentified group 1 capsules) (n=3117), the R3 outer core type was the most common (34.1%), followed by R1 (30.8%) and K-12 (14.6%), while R4 and R2 were less frequently observed (9.2% and 11.2%, respectively). Among specific K-antigens (Figure 5C), R1 was highly associated with K1, K15, K23, and K96, with frequencies ranging from 63% to 92%. By contrast, K52 and K98 showed the lowest representation of R1 (6.1% and 7.7%). The R4 outer core type was predominantly linked to K4 and K98 (61.5% each), while K52 was most strongly associated with R3 (67.3%).

These findings, showing LPS outer core enrichment in specific capsule types and across broad capsule groups, suggest that the presence and type of capsule might be influenced by the distribution of LPS outer core types, with R1 being particularly enriched in capsule-containing strains. Conversely, the R3 LPS outer core type was more frequently associated with capsule-deficient strains. However, it was highly enriched in K52-type capsules, suggesting structural or functional advantages for the co-occurrence of the R3 LPS outer core with specific capsule structures.

## Discussion

Our study introduces a rapid and comprehensive framework for *in silico* and *in vitro* typing of *E. coli* R1, R2, R3, R4, and K-12 LPS outer core structures by targeting unique regions within the *waa* locus. Using nine novel oligos, we achieved robust typing with a 99% success rate *in silico* across 4549 complete *E. coli* genomes from the NCBI database using our LPSTyper tool. We successfully validated these new improved oligos (Table 1) on reference strains using a simple *in vitro* PCR method (Table 2 and Figure 2).

### LPS outer core type patterns in commensal and pathogenic E. coli

Our analysis of LPS outer core types in 4549 genomes, including commensal and pathogenic strains, revealed distinct distribution patterns. The R1 outer core type emerged as the most prevalent, as reported in previous works [13, 14, 41, 42], followed by R3, K-12, and R4, with R2 being the least common. In earlier studies, R1 type was observed in 68.3% of 180 strains typed using mouse monoclonal antibodies [13]; however, nearly 30 years later, it was detected in only 33.9% of 499 strains using PCR typing [42]. Our current study of 4549 strains identified the R1 type in 35.5% of *E. coli* strains (Figure 3A), indicating that while R1 remains a prominent core type, its relative abundance diminishes in larger, more diverse genomic datasets. Our data also reveal a significant enrichment of the R1 LPS core type in specific pathogenic *E. coli* strains, including UPEC (59.7%), NMEC (92.6%), ETEC (65.9%), and APEC (68.6%). The prevalence of the R1 core structure may confer a selective advantage by facilitating the acquisition of virulence genes, enhancing colonisation in various host tissues, or improving survival and replication in the circulatory system [14, 43]. This hypothesis warrants further investigation to fully elucidate the role of the R1 core in the biology and pathogenesis of *E. coli*.

### Exploring the connection between O-antigens and LPS outer core types in E. coli

The relationship between serotypes and LPS outer core structures in *E. coli* remains a subject of ongoing investigation, as the O-antigen is covalently attached to a specific residue of the LPS outer core[44]. Previous studies on smaller *E. coli* genomic datasets lacked sufficient sample sizes to determine statistical significance or identify associations between these features [14, 42]. However, our data reveal significant associations between specific O-antigen types and LPS outer core types (Figure 5A). Notable examples identified in this work include K-12 with O16 and O25 serotypes, R2 with O89, R3 with O157-antigen, and R4 showing moderate associations with O102 and O86 serotypes. We also observed that while LPS outer core types can be linked to multiple O-antigen serotypes, each O-antigen type generally exhibits a preference for a single LPS outer core type. However, some O-antigens such as O1, O5, O83, O128, O141, and others, demonstrated compatibility with multiple core types (Figure 5A and Table S5). This ability to link diverse O-antigens to each LPS outer core type underscores the dynamic nature of LPS biosynthesis and likely relies on the precise enzymatic transfer of the O-antigen from a pyrophosphate-undecaprenyl carrier to the LPS core by WaaL, an enzyme with broad donor substrate specificity [45]. Further investigation is needed to understand this variability in O-antigen-core linkages fully.

Moreover, particular combinations of O-antigen and outer core type may play a role in enhancing bacterial immune evasion and host adaptation. For instance, enterohemorrhagic *E. coli* (EHEC) strains are predominantly characterised by the R3 outer core type and the O157 serotype, which account for 78.5% of the phylogenetic group E in our data. The prevalence of the R3 and O157 combination suggests there is a selective advantage for pathogenicity, host colonisation, or defence against phage-mediated attack. Beyond bacteriophage resistance, evidence indicates that predatory protozoa, such as *Acanthamoeba castellanii,* can influence *E. coli* survival by selectively modulating bacterial consumption based on *E. coli* LPS outer core type and O-antigen composition [46, 47]. This suggests that variations in LPS structure may indirectly enhance virulence and contribute to environmental persistence by reducing susceptibility to predation.

### Hybrid and Novel LPS Outer Core Structures

In this work, we identified novel non-canonical *waa* locus gene structures (Table S11), including genomic rearrangements between R2 and K-12 LPS outer cores, gene duplications, and the presence of transposable elements, highlighting the dynamic nature of the *waa* locus (Figure1). These findings align with previous studies suggesting recombination and horizontal gene transfer drive LPS variability, potentially influencing host immune evasion and bacterial environmental adaptability [48–50].

Notably, we show that the R2 outer core type, the rarest *E. coli* LPS type in our dataset, was associated with several hybrid variations involving the K-12 core type (Figure 1 and Table S11). This association between R2 and K-12 outer core types adds evidence to the notion that the R2 outer core type itself is a hybrid of *E. coli* K-12 and *Salmonella enterica* LPS core types [51]. In the K-12/R2 gene arrangements we identified, eight K-12 *E. coli* strains harboured *waaK* genes instead of *waaU*, whereas one R2 strain contained *waaU* instead of *waaK* (Table S11). The *waaK* gene encodes UDP-N-acetylglucosamine:(glucosyl) lipopolysaccharide α1,2-N-acetylglucosaminyltransferase, which in the R2 outer core plays a critical role by adding a terminal α1,2-linked GlcNAc side branch [51]. This modification is essential for recognition by the O-polysaccharide ligase WaaL. By contrast, in the K-12 outer core, WaaU, an ADP-L-glycero-β-D-manno-heptose:LPS heptosyltransferase IV, attaches a fourth heptose residue to the third glucose, serving as the site for O-antigen attachment. At first glance, these hybrid structures might be expected to resemble either the K-12 or R2 core. However, additional modifications introduced by WaaS and WabA could result in structures that do not precisely match either parental type. Notably, the hybrid K-12/R2 outer cores can accommodate O-antigens not supported by the original K-12 structure, such as O127, O19, and O58, suggesting that the structural flexibility of WaaL is influenced by the modified core. Interestingly, the single R2/K-12 hybrid carried the O3 antigen (Figure 1, Table S11), which is typically associated with R1-R4 outer core types, and no additional O-antigen structures unrelated to R2 were incorporated into this chimera. Due to the limited number of hybrid strains identified, it remains unclear whether specific O-antigens preferentially associate with these hybrid LPS outer cores or whether additional structural constraints influence their compatibility. Further studies incorporating a broader genomic dataset would be necessary to clarify these relationships.

Additionally, our study identified an R1/R4 hybrid in the ECOR collection strain ECOR46. The formation of such a hybrid core is thought to require the co-expression of *waaL* genes specific to R1 and R4, along with *waaV* and *waaX* [14]. Notably, ECOR46 meets these criteria, possessing R1- and R4-specific *waaL*, *waaV* from R1, and *waaX* from R4. This result suggests ECOR46 should not be strictly classified as R1 or R4. However, the exact structure of its LPS chimera remains unclear, requiring further experimental characterisation.

In conclusion, this study underscores the evolutionary plasticity of the *waa* locus and highlights the effectiveness of our primers and *in silico* typing tool in revealing the diversity and distribution of LPS outer core structures in *E. coli*. The strong associations between LPS types, pathogenicity, phylogenetic groups, and genomic rearrangements emphasise the significance of LPS variability in bacterial fitness, adaptation, and virulence. Our findings demonstrate that the *in silico* typing method is the most reliable approach for genome-based analysis and should be prioritised when genomic sequences are available. Moreover, the differences between this work and prior reports [14, 42] highlight the need for regular updates to typing tools to accommodate novel genomic variability uncovered by increasing deep sequencing of the biosphere. As more genomic sequences become available, expanding analyses to encompass incomplete genomes will likely reveal a broader diversity of hybrid LPS outer core types. These efforts will strengthen our understanding of LPS structural evolution and its role in *E. coli* ecology and pathogenicity.

## Acknowledgements

We recognise that this research was conducted on the traditional lands of the Wallumattagal clan of the Dharug nation. We thank Prof. Kirill Alexandrov (QUT) for providing laboratory space that facilitated some of the work.

## Author Contributions

PRJ: Funding acquisition, Project administration, Supervision, Writing – review & editing.

RW: Investigation, Writing - review & editing.

ET: Conceptualisation, Data curation, Formal analysis, Investigation, Methodology, Software, Validation, Visualisation, Writing - original draft, Writing - review & editing.

